# New insights on malaria parasite adaptation yielded by samples archived for more than 50 years

**DOI:** 10.1101/2025.03.11.642566

**Authors:** Alfred Amambua-Ngwa, Mouhamadou Fadel Diop, Christopher J. Drakeley, Umberto d’Alessandro, Dominic P. Kwiatkowski, David J. Conway

## Abstract

Understanding evolution of human pathogens requires looking beyond effects of recent interventions or known epidemiological changes. To study malaria parasites prior to widespread drug selection, *Plasmodium falciparum* genome sequences were analysed from the oldest set of archived research samples yet identified, placental blood collected in the Gambia between 1966 and 1971. Genomic complexity within infections was high, infections were genetically unrelated, and no drug resistance alleles were detected. Strong signatures of positive selection are clearly seen at multiple loci throughout the genome, most of which encode surface proteins that bind erythrocytes and are targets of acquired antibody responses. Comparison of population samples obtained over a following period of almost 50 years revealed major directional allele frequency changes at several loci apart from drug resistance genes. Exceptional changes over this time are seen at *gdv1* that regulates the rate of parasite sexual conversion required for transmission, and at the unlinked *Pfsa1* and *Pfsa3* loci previously associated with infection of individuals with sickle-cell trait. Other affected loci encode surface and transporter proteins warranting targeted functional analyses. This identification of key long-term adaptions that have not reached equilibrium is important for understanding and potentially managing future evolution of malaria parasites.

## Introduction

Population genomic studies of malaria parasites have yielded important insights on selection operating under different malaria control contexts. These have helped understand drug resistance emergence and spread, to inform antimalarial treatment and chemoprevention policies ^1-7^. However, it is important to understand also the modes of selection that operated prior to drug resistance emergence, and which continue to affect interactions between parasites and their hosts ^8^. Ancient DNA from excavated remains of infected individuals have begun to give insights on past evolution of malaria parasites ^9^, and museum specimens may also contain parasite DNA for analysis ^10,11^, but such sources have yielded only sporadic positive samples so far.

Early research archives with sufficient material to enable population-based analyses on historic selective processes are particularly valuable. Previous analyses of archived population samples of *Plasmodium falciparum* taken at different times in The Gambia dating back to the 1980s have revealed the local history of drug selection ^1,12-14^. Chloroquine resistance first became common due to selection for variants of the *aat1* and *crt* drug transporter genes, following which sulphadoxine-pyrimethamine was added as first line treatment and antifolate resistance became common due to selection of *dhfr* and *dhps* variants encoding altered target sites ^1,14^. Artemisininin-based combination therapy was introduced in 2008, although malaria incidence in The Gambia started declining earlier ^15-19^. Besides drug resistance, genomic studies on parasites in The Gambia have highlighted the operation of balancing selection on targets of immunity ^20^, and allowed comparison of local signatures of selection with those occurring in other parasite populations in Africa ^13,21,22^.

Aiming to understand selection on malaria parasites in the more distant past, and over a longer period, records of earlier studies were examined to seek the oldest archived samples containing parasite material. These were found to be placental blood samples collected in The Gambia in 1966-1971 which originally enabled the first analysis of *P. falciparum* isoenzyme variation ^23^ and description of polymorphic antigens ^24,25^, undertaken in the era before methods to analyse DNA were available. Here, the parasite genome sequences from the entire available set of samples are analysed, revealing the key features of positive selection that had been operating on parasites long before the detection of any drug resistance. More recent samples from the same area were then added for the longest-term temporal analysis yet performed, which identified new targets of selection occurring over the subsequent period of almost 50 years. This identifies ways in which parasites have continued to evolve and adapt to their human hosts, highlighting targets for development of future tools to sustain malaria control in future.

## Results

### Identifying targets of selection on malaria parasites prior to drug resistance

Archived *P. falciparum*-positive placental blood samples collected in 1966-1971 from the coastal area of The Gambia were identified and processed for parasite genome sequencing. Fifty-four (90%) of 60 samples yielded >80% genome-wide coverage at a read depth of at least 5x (Supplementary Table 1), enabling a powerful analysis of genome-wide diversity.

Initially, the genome sequences of the 54 infections were considered together as a population sample from this era, enabling analysis of diversity throughout the genome. To analyse population-wide *P. falciparum* allele frequencies for all SNPs, for each infection the predominant genotype profile for each SNP was defined by the majority of sequence reads mapped, and genome-wide tests for positive selection were applied to the profiles from all infections. These data show that strong positive selection was operating on multiple loci in the parasite genome long before drug resistance emerged in this population, and there were no drug resistance loci under selection (Fig. 1). Twenty-six genes in sixteen separate genomic regions had the highest values of the beta score index which is generally an indication of long-term balancing selection (Fig. 1 and Table 1), and 14 of these genes contained SNPs with significant integrated haplotype scores (iHS), indicating recent positive directional selection on some alleles (Fig. 1 and Table 1).

**Fig. 1.**
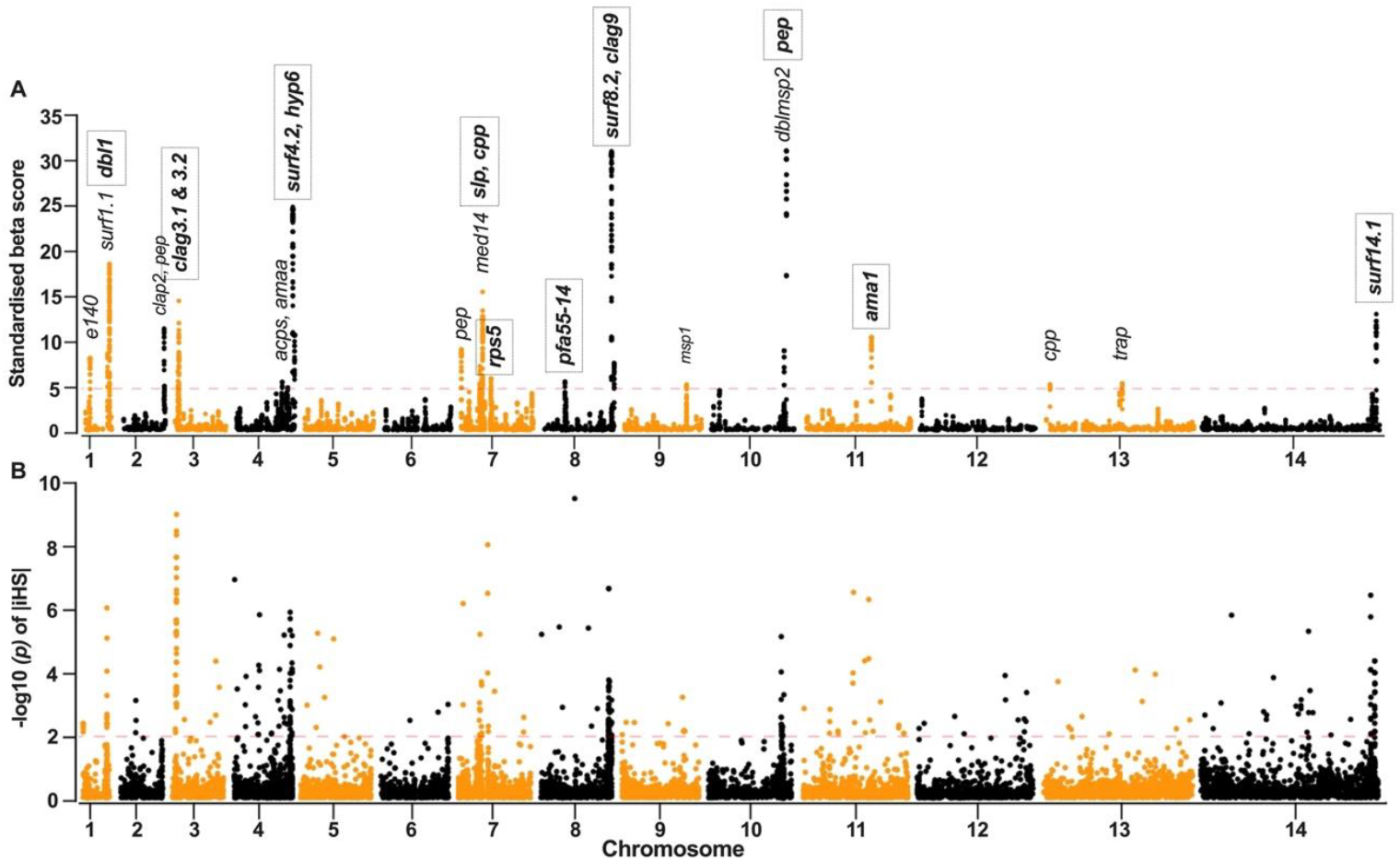
Genome-wide scans for signatures of selection on the *P. falciparum* population in The Gambia sampled in 1966-1971, many years prior to emergence of drug resistance. **A**. The beta score index across sliding windows of SNPs for all 14 chromosomes identifies loci with correlated SNP allele frequency distributions indicating balancing selection. Twenty-six genes in the windows with the highest beta scores are labelled, and 14 of these (presented in bold font within boxes and in Table 1) also contained at least one SNP with a significant integrated haplotype score (iHS). **B**. The -log10 p value for the integrated haplotype score (iHS) which indicates recent directional selection on the basis of extended haplotype homozygosity is shown for each SNP. Most of the genes showing these signatures of positive selection encode parasite surface proteins that interact with host cells and are targets of antibody responses. Values for beta score and iHS indices of all genes analysed genome-wide are listed in Supplementary Tables 2 and 3.

**Table 1.**
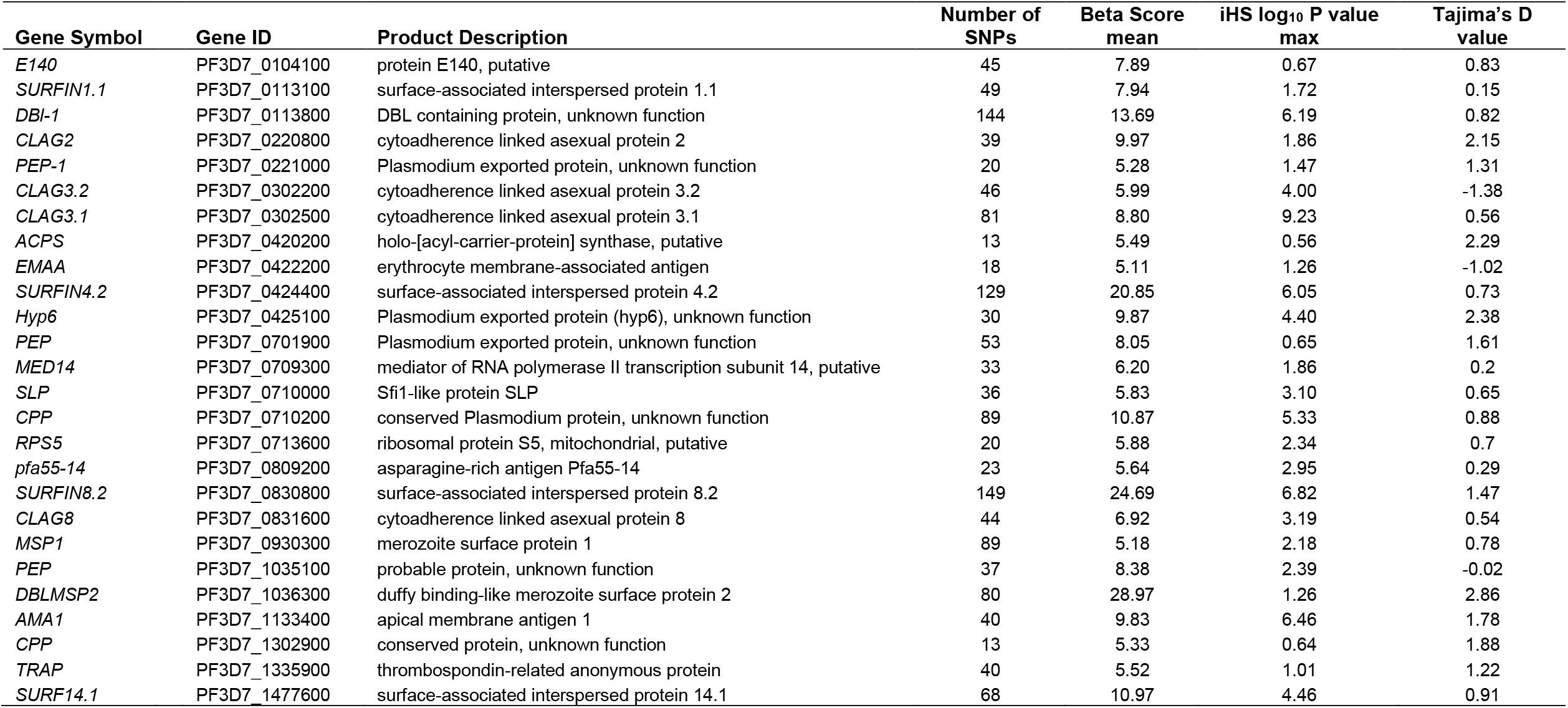
Population-based genome-wide scan of 54 archived *P. falciparum* infection samples from 1966-1971 identifies 26 genes with strong signatures of positive selection in an era prior to drug resistance. These are indicated by the mean beta score for each gene, listed alongside significance of integrated haplotype score (iHS) and Tajima’s D.

| Gene Symbol | Gene ID | Product Description | Number of SNPs | Beta Score mean | iHS log<sub>10</sub> P value max | Tajima's D value |
| --- | --- | --- | --- | --- | --- | --- |
| *E140* | PF3D7_0104100 | protein E140, putative | 45 | 7.89 | 0.67 | 0.83 |
| *SURFIN1.1* | PF3D7_0113100 | surface-associated interspersed protein 1.1 | 49 | 7.94 | 1.72 | 0.15 |
| *DBI-1* | PF3D7_0113800 | DBL containing protein, unknown function | 144 | 13.69 | 6.19 | 0.82 |
| *CLAG2* | PF3D7_0220800 | cytoadherence linked asexual protein 2 | 39 | 9.97 | 1.86 | 2.15 |
| *PEP-1* | PF3D7_0221000 | Plasmodium exported protein, unknown function | 20 | 5.28 | 1.47 | 1.31 |
| *CLAG3.2* | PF3D7_0302200 | cytoadherence linked asexual protein 3.2 | 46 | 5.99 | 4.00 | -1.38 |
| *CLAG3.1* | PF3D7_0302500 | cytoadherence linked asexual protein 3.1 | 81 | 8.80 | 9.23 | 0.56 |
| *ACPS* | PF3D7_0420200 | holo-[acyl-carrier-protein] synthase, putative | 13 | 5.49 | 0.56 | 2.29 |
| *EMAA* | PF3D7_0422200 | erythrocyte membrane-associated antigen | 18 | 5.11 | 1.26 | -1.02 |
| *SURFIN4.2* | PF3D7_0424400 | surface-associated interspersed protein 4.2 | 129 | 20.85 | 6.05 | 0.73 |
| *Hyp6* | PF3D7_0425100 | Plasmodium exported protein (hyp6), unknown function | 30 | 9.87 | 4.40 | 2.38 |
| *PEP* | PF3D7_0701900 | Plasmodium exported protein, unknown function | 53 | 8.05 | 0.65 | 1.61 |
| *MED14* | PF3D7_0709300 | mediator of RNA polymerase II transcription subunit 14, putative | 33 | 6.20 | 1.86 | 0.2 |
| *SLP* | PF3D7_0710000 | Sfi1-like protein SLP | 36 | 5.83 | 3.10 | 0.65 |
| *CPP* | PF3D7_0710200 | conserved Plasmodium protein, unknown function | 89 | 10.87 | 5.33 | 0.88 |
| *RPS5* | PF3D7_0713600 | ribosomal protein S5, mitochondrial, putative | 20 | 5.88 | 2.34 | 0.7 |
| *pfa55-14* | PF3D7_0809200 | asparagine-rich antigen Pfa55-14 | 23 | 5.64 | 2.95 | 0.29 |
| *SURFIN8.2* | PF3D7_0830800 | surface-associated interspersed protein 8.2 | 149 | 24.69 | 6.82 | 1.47 |
| *CLAG8* | PF3D7_0831600 | cytoadherence linked asexual protein 8 | 44 | 6.92 | 3.19 | 0.54 |
| *MSP1* | PF3D7_0930300 | merozoite surface protein 1 | 89 | 5.18 | 2.18 | 0.78 |
| *PEP* | PF3D7_1035100 | probable protein, unknown function | 37 | 8.38 | 2.39 | -0.02 |
| *DBLMSP2* | PF3D7_1036300 | duffy binding-like merozoite surface protein 2 | 80 | 28.97 | 1.26 | 2.86 |
| *AMA1* | PF3D7_1133400 | apical membrane antigen 1 | 40 | 9.83 | 6.46 | 1.78 |
| *CPP* | PF3D7_1302900 | conserved protein, unknown function | 13 | 5.33 | 0.64 | 1.88 |
| *TRAP* | PF3D7_1335900 | thrombospondin-related anonymous protein | 40 | 5.52 | 1.01 | 1.22 |
| *SURF14.1* | PF3D7_1477600 | surface-associated interspersed protein 14.1 | 68 | 10.97 | 4.46 | 0.91 |

Most of these parasite genes under positive selection encode surface proteins that are targeted by immune responses. These include merozoite surface proteins (MSP1, MSPDBL2), four members of one family of proteins expressed on merozoites or infected erythrocytes (Surfins 1.1, 4.2, 8.2 and 14.1), three members of another family of surface and rhoptry apical organelle proteins (CLAGs 2, 3.1/3.2, and 8), micronemal apical organelle proteins involved in erythrocyte invasion (EBA140 and AMA1) and a sporozoite surface protein (TRAP). Others encode a less characterised protein with a Duffy binding-like domain (DBL1), as well as uncharacterised proteins predicted to be exported from the intraerythrocytic parasite. Most genes with these signatures of selection are in or close to chromosomal subtelomeric regions, and are heterochromatic loci that show epigenetic variation in expression ^26^.

Using another approach to identify genes with alleles at more intermediate frequencies than expected, a scan of Tajima’s D values was performed on the genome-wide data from the 1966-1971 population (Supplementary Table 4, Supplementary Fig. 1). Twenty three of the 26 genes with highest beta scores also had positive values of Tajima’s D (Table 1 and Supplementary Fig. 1). This is consistent with strong balancing selection operating on these loci. In many cases this is likely to be due to frequency-dependent selection exerted by the strong memory component of human B and T cell responses, as previously noted for known targets of immunity ^20,27,28^.

### Genomic complexity of infections compared to more recent population samples

Many of the individual infection samples from 1966-1971 contained significant *P. falciparum* genomic sequence diversity, indicating that these contained multiple haploid parasite genotypes. The within-infection fixation index (*F*_WS_ which has a value of less than 1.0 when more than one genotype is present) had values of < 0.95 for 38 (70%) of the 54 isolates (Fig. 2A, Supplementary Table 5), indicating that most infections contained multiple different parasite genotypes. The overall mean *F*_WS_ value was 0.76, much lower than 1.0 and driven by low values from infections that contained very high levels of parasite genomic diversity. In more recent population samples from across Africa, such profiles have been seen only in areas with intense malaria transmission, whereas in moderate to low transmission areas within-infection diversity is much lower ^29,30^.

**Fig. 2.**
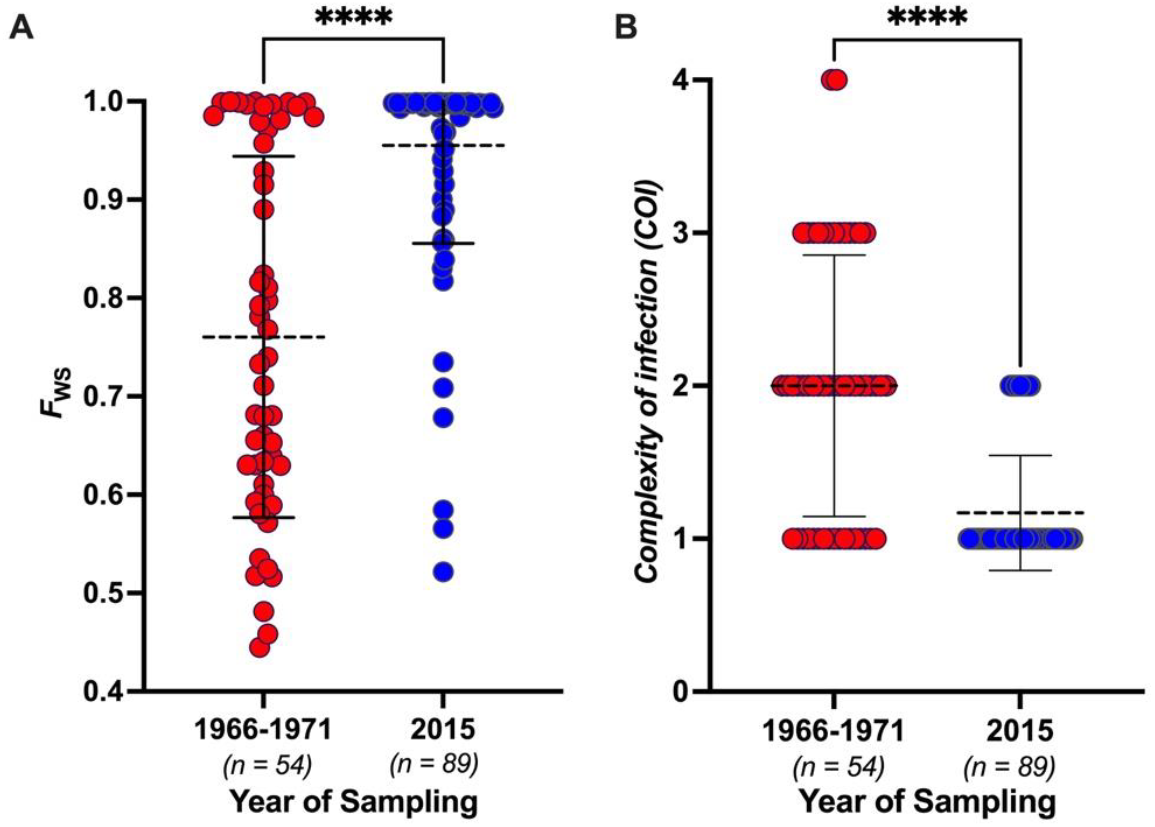
Genomic complexity within *Plasmodium falciparum* infections sampled in The Gambia in 1966–1971 and in 2015 (shown in red and blue points respectively). **A**. Within-infection fixation index (*F*_WS_) which is an inverse measure of genomic complexity (values of 1.0 corresponding to single-genome infections), with each infection represented by a separate point. Mean values for each sampling period are shown by dashed horizontal lines. **B**. ‘Complexity of Infection’ (COI) estimations using the REAL McCOIL method. The differences between sampling periods were highly significant (P<0.0001), indicated by asterisks ****. The values of *F*_WS_and COI for the individual infection samples in 1966-71 and 2015 are listed in Supplementary Tables 5 and 7 respectively.

To analyse the genomic diversity of infections from the same coastal area of The Gambia almost five decades later, parasite genomic DNA was extracted from blood samples taken from patients with uncomplicated malaria attending Brikama Hospital in 2015, and sequenced for analysis using the same methods used for the oldest archived samples (Supplementary Table 6). Within-infection diversity among 89 samples collected in 2015 was much lower, with most *F*_WS_ values close to 1.0 and an overall mean *F*_WS_ value of 0.96 (Mann-Whitney test comparing the sampling periods, P < *0*.*0001*) (Fig. 2A, Supplementary Table 7). Using a different approach to estimate ‘Complexity of Infection’ (COI) as the minimum number of distinct genotypes detectable in each sample, the levels of mixedness was much lower in 2015 than in 1966-1971 (Fig. 2B, Supplementary Table 7)(P *< 0*.*0001*).

### Population-wide genomic diversity compared to more recent population samples

To further compare the population samples, the extent of genomic differences among different infections was analysed. For this purpose, the dominant *P. falciparum* profile for each infection was considered, represented by SNP variants in the majority of mapped sequence reads. The population-wide diversity between infections was high among the archived samples from 1966-1971, and also among the more recent samples from 2015 (Fig. 3). Visualisation of overall diversity patterns by multidimensional scaling analysis revealed slightly more population structure among the 2015 samples, including a minority of infections being outliers (Fig. 3A). Hierarchical clustering of pairwise similarity between the genomic profiles shows a high diversity, as expected from recombination resulting from sexual reproduction that occurs during transmission by mosquitoes (Fig. 3B). A minority of infections showed inter-relatedness, although this was particularly rare among the older samples, as only two pairs of samples from 1966-1971 showed >95% identity by state (IBS) whereas there were 26 such related pairs in 2015.

**Fig. 3.**
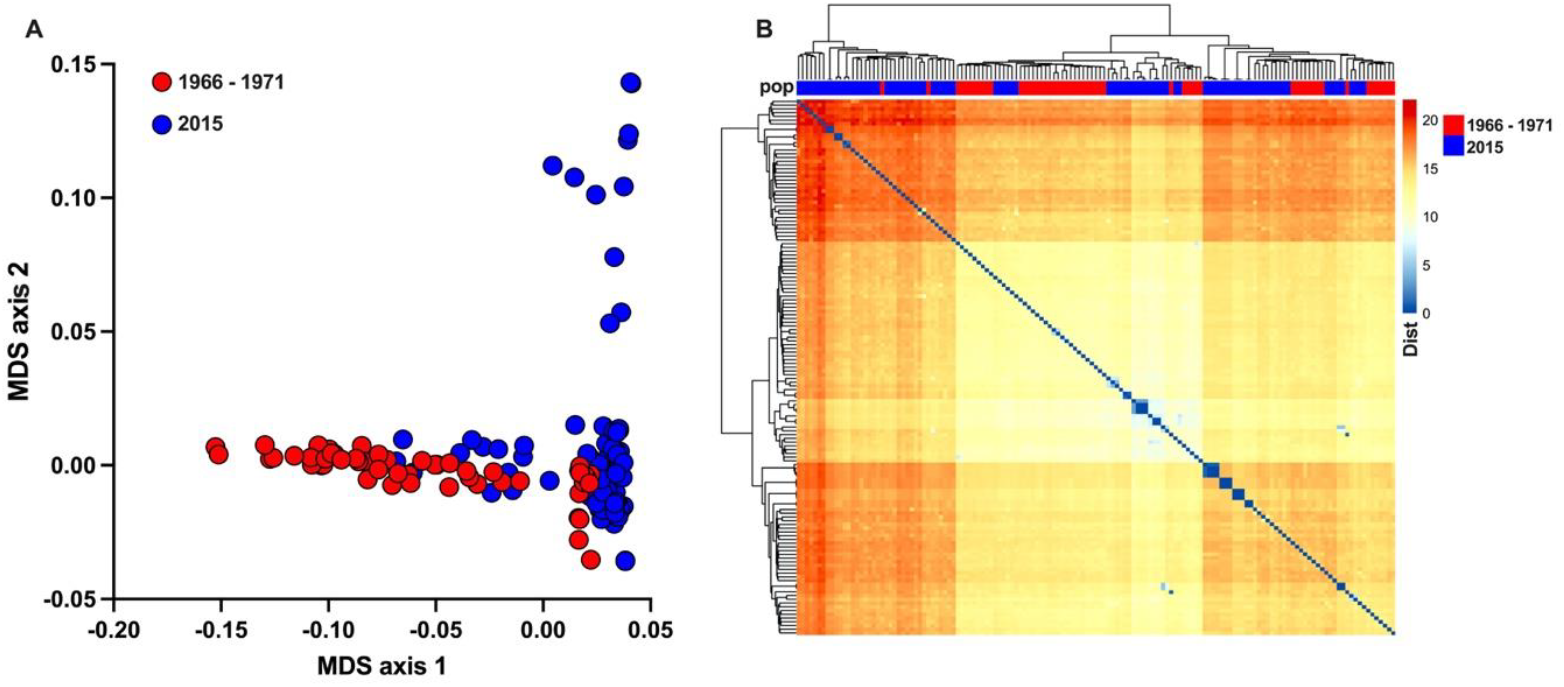
Population-wide diversity of *P. falciparum* genomic profiles in The Gambia in 1966-71 (shown in red) and 2015 (blue). **A**. A wide range of between-infection genomic diversity shown as a scatter plot of axes 1 and 2 of multidimensional scaling (MDS) based on the genomic profile of each of the infection samples, considering the predominant SNP at each locus within each individual infection. **B**. Neighbour-Joining dendrogram clustering and heatmap based on the genome-wide pairwise SNP differences between different infections. Profiles from of the different eras were overlapping and interspersed. Most infections were unrelated, although a minority of infections showing high relatedness were identified in the more recent population sampled in 2015, identifiable by larger blue squares along the diagonal of the matrix heatmap.

### Identifying parasite loci with exceptional allele frequency changes over time

To identify which regions of the *P. falciparum* genome have undergone the most pronounced allele frequency changes since 1966-1971, a comparison was performed with the 2015 data (Fig. 4A, Supplementary Table 8). By applying the temporal *F*_ST_ fixation index, it is clear that discrete loci have undergone marked changes in frequency, whereas the genome-wide background shows minimal frequency change. Four of the top ten genomic loci with high fixation indices contain drug resistance genes (*dhfr, aat1, crt*, and *dhps*) already known to have been under selection in The Gambia ^1,14^. There were no drug resistance alleles at any of these loci in the 1966-71 samples, but these became common from the 1980s onwards (Supplementary Table 9, Supplementary Fig. 2) ^1,14^. Different processes of selection have operated on the other genomic loci showing exceptional levels of allele frequency change over time (*pfsa1, gdv1, pfsa3, msp7, mfs6* and *cpp-5*). Analysis of allele frequencies in additional population samples collected at intervening time points in The Gambia allow the temporal trajectories to be more fully understood (Fig 4B, Supplementary Fig. 3 and Supplementary Table 9).

**Fig. 4.**
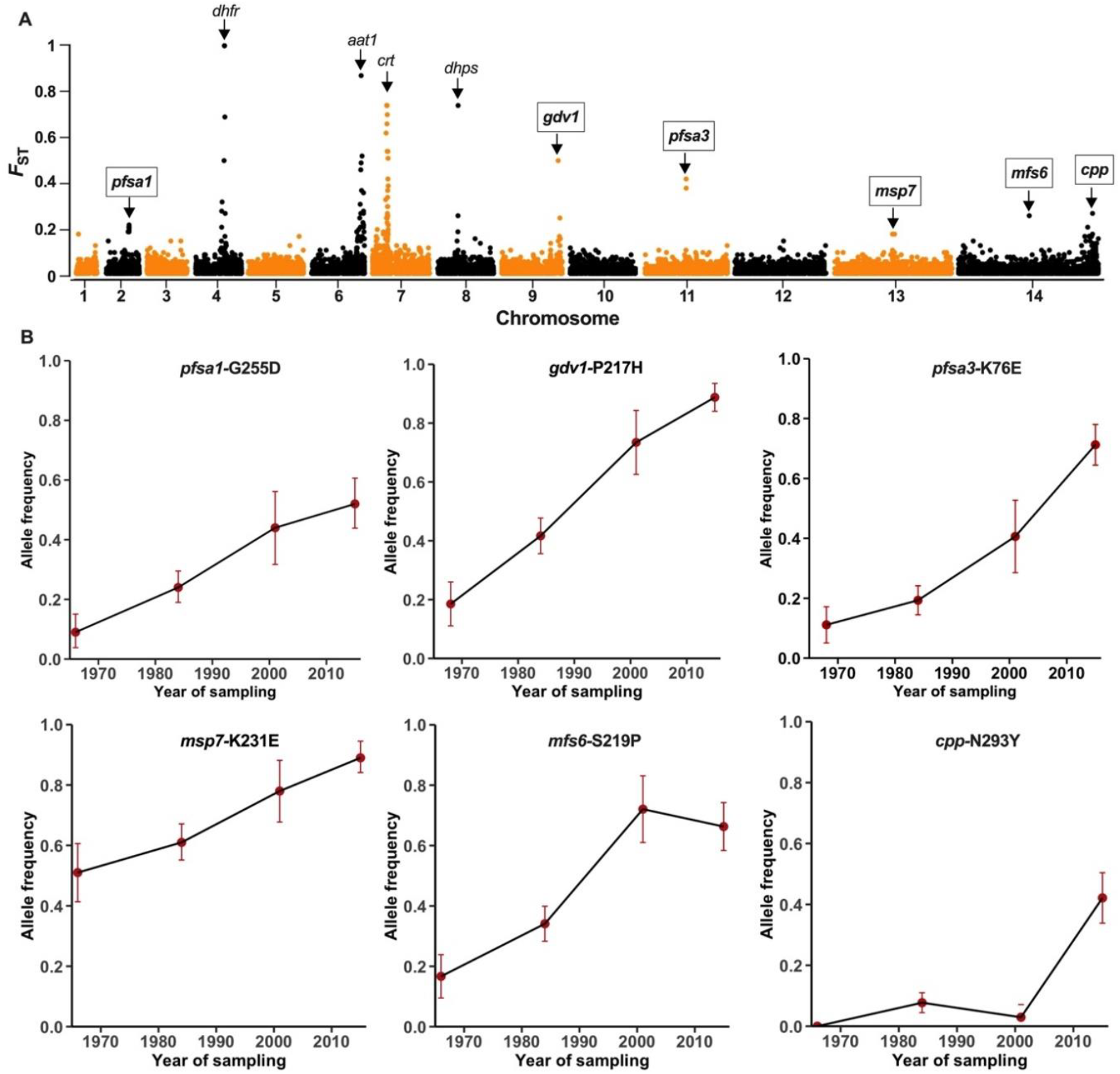
*Plasmodium falciparum* genomic polymorphisms with the most pronounced changes in allele frequencies over a period of almost 50 years in The Gambia. **A**. Genome-wide scan for temporal differentiation shown as *F*_ST_ temporal fixation indices on SNP allele frequencies in the oldest population samples (N = 54 from 1966-71) compared with more recent population samples also sequenced in this study (N = 89 from 2015). The labels identify the top ten genomic loci showing high temporal allele frequency differentiation. Four of these loci (with unboxed labels) are known drug resistance genes previously shown to be under long-term selection in The Gambia (*dhfr, aat1, crt*, and *dhps*) ^1,14^ for which resistance alleles had zero frequency in the 1966-71 samples. The other six loci (with boxed labels) showing exceptional frequency change over time are unrelated to drug resistance. **B**. Trajectories of temporal allele frequency changes at the most differentiated SNPs in these six loci are shown by also including previously obtained parasite genome sequence data ^1^ from other intervening times (1984 and 2001), alongside the data from 1966-71 (plotted here using the median sample year 1968) and 2015. The *Pfsa1* and *Pfsa3* loci are associated with infection of individuals with sickle cell trait ^8^; *gdv1* regulates parasite sexual commitment ^32^ and expression of a major variant merozoite surface protein ^33^; *msp7* encodes a merozoite surface protein ^65^; *mfs6* encodes a membrane transporter ^39^; *cpp* encodes a conserved *Plasmodium* protein with gene ontology prediction as an intergral membrane component.

Besides the drug resistance-associated loci, the gene with the highest temporal differentiation is *gdv1* (*PF3D7_0935400* on chr 9) that regulates parasite sexual commitment ^31,32^ and controls expression of the MSPDBL2 antigen in mature schizonts ^33^. The most temporally-differentiated polymorphism in the *gdv1* gene (SNP causing the codon change P217H), involved the 217H allele increasing in frequency from less than 20% to almost 90% (Fig. 4B). The temporal data here showed allele frequency change at some closely-linked loci (Supplementary Fig. 3 and Supplementary Table 9), including the *clag9* gene (*PF3D7_0935400*, Supplementary Fig. 4).

Notably, two of the other loci showing marked frequency change over time are likely to affect parasite replication in specific human hosts. Variation at the *Pfsa3* locus on chr 11 has been associated with parasite infection of individuals with sickle-cell trait (carrying the HbS allele that protects against severe malaria) ^8^. The most temporally differentiated polymorphism at this chromosomal locus is codon K76E in gene *PF3D7_1127000* (Fig. 4B) encoding a putative tyrosine phosphatase observed in the parasite food vacuole ^34^, while codon D7V polymorphism in the same gene shows almost the same level of temporal change (Supplementary Fig. 3). Remarkably, the *Pfsa1* locus on chr 2 has also been associated with infections of individuals having sickle-cell trait, and appears to be under parallel selection, showing strong linkage disequilibrium with *Pfsa3* although it is physically unlinked ^8^. At the core of the *Pfsa1* locus are six polymorphic codons in the Acyl-CoA synthetase gene (*PF3D7_0215300*) showing temporal differentiation at approximately the same level (Fig. 4 and Supplementary Fig. 3).

Another locus with high temporal differentiation maps to the *msp7* gene (*PF3D7_1335100* on chr 13) which encodes a merozoite surface protein, near genes encoding other surface antigens, including the sporozoite surface protein TRAP, and Rh2b involved in erythrocyte invasion which have previously shown significant signatures of selection ^35-37^ . The *trap* gene (*PF3D7_1335900*) contains a SNP encoding a D166N codon polymorphism that also shows significant temporal changes over the period analysed (Supplementary Table 9, Supplementary Fig. 4).

The *mfs6* gene on chr 14 encodes a major facilitator superfamily-domain-containing protein which is a membrane transporter ^38^. The orthologue of this protein in the rodent malaria parasite *P. berghei* is required for optimal replication of parasites during the blood stage of infection, and essential for enabling a new infection to be established from an infected mosquito bite ^39^. Experiments with *P. falciparum* clinical isolates would be needed to test whether the variants of this gene affect intrinsic multiplication rates, which vary significantly among isolates in The Gambia ^40^ and elsewhere. The other locus with significant temporal change located distally on chr 14 has only previously been denoted as a conserved *Plasmodium* protein (*cpp*) as it has orthologues in all species, but it is predicted by gene ontology to be an integral membrane protein.

## Discussion

Analysis of the oldest archived endemic population samples of malaria parasites here has revealed the principal targets of positive selection operating prior to drug resistance.

Furthermore, comparison with more recent samples has uncovered other adaptive changes occurring progressively over decades. Originally, the targets of selecton were mostly genes encoding parasite surface and apical organelle proteins that bind to erythrocytes, as well as proteins embedded in the infected erythrocyte membrane. It is notable that the major targets of natural selection on humans by malaria are genomic polymorphisms affecting erythrocytes ^41,42^. Results here indicate that balancing selection operated on most of the affected parasite genes, consistent with a frequency-dependent selective effect of naturally acquired immune responses. Higher malaria transmission and superinfection rates would likely have caused more intense immune selection than occurs today, and indeed more individual antigen genes were identified as under selection compared with more recent population samples in the same country ^12^.

The high level of within-infection *P. falciparum* diversity in the the oldest archived samples collected shows that malaria transmission and superinfection in The Gambia at the time was intense, consistent with epidemiological reports from that era ^18,43^. The samples were from placental blood, whereas more recent samples analysed were from peripheral blood of malaria cases, but previous studies of placental blood samples have not indicated greater *P. falciparum* within-infection diversity compared to peripheral blood samples ^44,45^, and the temporal change in within-infection diversity reflects the significant malaria reduction in malaria in the country ^15,16,19^.

Analysis of subsequent genomic changes in the parasite population over almost 50 years reveal ongoing adaptions in parasite-host interactions. As expected, drug resistance variants were completely absent from the old archived samples but appeared later in The Gambia and then rapidly increased in frequency, whereas most other significant allele frequency changes involve variants already present in the late 1960s. Among the genomic loci not associated with drug resistance, the strongest evidence of directional selection over time is seen in the gametocyte development gene *gdv1*. This should be considered alongside previous analyses indicating differential selection on this locus in different endemic areas.

Notably, the polymorphism with highest temporal change here (SNP causing the codon polymorphism P217H) was previously shown to have the highest geographical differentiation among West African populations ^22,46^. The *gdv1* allele 217H that has become frequent in The Gambia is still relatively rare in most other West African populations (except for Senegal which closely neighbours The Gambia), indicating that the selection has operated within this particular area of West Africa ^46^.

Selection on *gdv1* is likely to reflect adaptation to altered transmission conditions, given its key role of in regulating parasite sexual commitment to gametocyte development ^32^. Interestingly, in Ghana where the *gdv1* 217H allele is much less common than in The Gambia ^46^, it has been reported to be associated with parasites having higher rates of gametocyte conversion in *ex vivo* culture ^47^. Interpretation and further testing requires recognition that the SNP at codon 217 has been shown to be in linkage disequilibrium with other polymorphisms in the coding sequence and in the 3’-intergenic region, as well as several neighbouring genes ^46^ including the *clag9* gene that encodes a protein with evidence of function in merozoites ^48^ and infected erythrocytes ^49^. Furthermore, the *gdv1* gene also regulates epigenetic expression of the variant merozoite surface antigen MSPDBL2 that is not involved in sexual commitment ^33^, so identifying the adaptive function of *gdv1* polymorphism may require consideration of more complex phenotypes.

It is remarkable that variants in both the *Pfsa1* and *Pfsa3* loci on chromosomes 2 and 9 respectively, associated with infections of sickle-cell trait individuals ^8^, show exceptional changes in allele frequencies over time. Previous analysis of *Pfsa3* genotype frequencies in Gambian samples from malaria patients indicated changes in allele frequency between 1999 and 2008 ^8^. Our results here show that changes have been more profound and continuous over a much longer period. As the frequency of sickle cell genotypes have not changed significantly in the local human population during this period, we suspect that these parasite variants affect fitness of infections in individuals with different levels of haemoglobin including those with anaemia. The mean levels of haemoglobin in children in The Gambia have increased substantially during the study period ^50,51^, partly due to a reduction of the malaria burden ^15,52^, and this may have had a selective effect on the parasite ^53^.

Such an approach of identifying and analysing archived biomedical samples can generally help understand historical and long-term selection. The potential will depend to some extent on original methods used for archiving and maintaining research specimens, distinct from approaches to the analysis of ancient DNA that may be obtained in smaller amounts from archeological specimens ^54,55^. For plant pathogens, this has usually involved dried herbarium samples, which has allowed informative analysis of plant pathogens back to the 19^th^ century ^56,57^. For human and animal pathogens, museum samples might contain some tissue material collected in an era prior to biomedical research on pathogens ^10,11^, although the numbers of such specimens are likely to be too low for analyses of population allele frequencies. The value of preserving research samples for pathogen analysis in the long-term future is thereby highlighted. As this study has uncovered key processes of selection on a malaria parasite species that still causes over half a million deaths annualy, it illustrates the importance of investing in sample maintenance by institutions in low income countries, as one pillar towards building more equitable research power ^58^.

## Online Methods

### Identification of P. falciparum-positive archived blood samples and DNA extraction

A search was undertaken for parasite population samples that might be considerably older than those previously sequenced (the oldest samples previously analysed were from 1984)^1^. Examination of archived materials from studies at the MRC Unit in The Gambia identified 61 *P. falciparum*-positive lyophilised placental blood samples from donors sampled in 1966-1971. As these samples had been originally analysed in the early 1970s in a study of parasite enzyme electrophoretic variation ^23^, it was anticipated that most would contain sufficient parasite material to enable selective whole genome amplification for Illumina paired-end short read sequencing. Under approval by the Joint Ethics Committee of the MRC Unit and the Gambian Government, all of these archived samples were processed for analysis by first extracting DNA from the lyophilised blood recovered from a single replicate tube for each sample. The lyophilised blood was effectively solubilised by addition to sample extraction buffer from a QIAamp® Blood Extraction Midi kit (QIAGEN, UK), and DNA was extracted using the same kit with the protocol as previously applied to samples of whole blood.

### Sequencing of P. falciparum genomes from archived blood samples

Whole genome sequencing was performed using the same methods as applied to recent samples in previous studies ^12^. Briefly, selective whole genome amplification to enrich for *P. falciparum* DNA compared to human DNA was performed using a previously described method ^59^, and paired-end short read (150bp) sequencing was performed on the Illumina HiSeq platform at the Wellcome Sanger Institute under the MalariaGEN pipeline, which generated parasite sequences for all except one of the 61 samples. The sequence data for all of these samples are available in the ENA database (accession numbers and data on read coverage are given in Supplementary Table S1). Variant calling followed established processes in the MalariaGEN Pf7 pipeline, with this analysis utilising biallelic SNP data with a minimum of 5 reads supporting each variant call within each infection sample, with MQ quality scores of at least 30. Following filtration to not more than 20% missingness and a population-wide minor allele frequency of at least 0.03, the analysis focused on 12,416 bi-allelic SNP variants genome-wide in 54 infection samples (the remaining six infection samples having much lower coverage were not analysed). To enable comparisons with parasites sampled from the same area of The Gambia almost 50 years later, Illumina short-read genome sequencing was performed on 89 *P. falciparum* infection samples from malaria cases presenting to Brikama Health Centre in 2015, and variant calling performed using the same method to allow analysis of the same set of genome-wide SNPs. Analyses of parasites sampled at different times representing intervening periods utilised sequence data from 1984 and 2001 that had been previously generated ^1,12,14,20^, with variant calling being performed in parallel with the other samples for the analysis here.

### Genomic complexity of infections and within-isolate fixation indices

Within each sample, the multiplicity of genomic backgrounds was determined using the within-infection fixation index Fws and an estimate of complexity of infection (COI). Fws was determined using the moimix R package ^60^ by analysis of 3927 SNPs with no data missingness in any infection sample and a minor allele frequency of at least 5% in both the 1966-71 and 2015 data. A measure of genomic complexity of infection (COI) that aims to enumerate clearly different genotypes was also generated using the REAL McCOIL method ^61^ with McCOILR package scripts employed that are available in the github repository (mpb).

### Population structure and pairwise relatedness

Population genotype VCF files were converted to plink using VCFtools. SNPs with pairwise LD were pruned using an r^2^>0.1 as cutoff over a sliding window of 50 consecutive SNPs and window jumps of 10 SNPs along each chromosome. From this, 6267 variants retained were used to generate pairwise identity by state (IBS) and genetic distance (1-IBS) matrix between all pairs of isolates. The distance matrix was used for dimension reduction in plink. Eigenvectors were plotted as scatter plots in R to visualise clusters.

### Identifying genomic regions under selection prior to drug resistance

To detect signatures of balancing selection in each population sample we first used the allele frequency based summary statistics, beta score (β), which detects ancient balancing selection followed by Tajima’s D, using all variants in non-repeat coding regions for genes with at least 3 coding polymorphic SNPs irrespective of minor allele frequency. The beta statistic was calculate using the python package BetaScan, which detects clusters of alleles at similar frequencies around a balanced locus as previously described ^62^. For this, the folded SNP frequencies were determined using VCFtools and fed to BetaScan with a -fold flag, retaining positive values. Tajima’s D was calculated across each gene using custom R scripts as previously described ^20^. We next applied the integrated haplotype score (iHS) statistic, for detecting evidence of recent positive directional selection from extended haplotype homozygisty calculated with the Rehh R package ^63^.

### Identifying P. falciparum genomic changes occurring in The Gambia after 1966-1971

To identify genomic regions differentiated between 1966-1971 and 2015 in the Gambia, 12416 common biallelic SNPs were employed to conduct a scan using ‘diff_stat’ function with mmod package in R. The *F*_*S*T_ index was calculated for each SNP and visualised on a genome-wide Manhattan plot using package qqman in R. Regions of SNPs in genes with highlty differentiating missense (non-synonymous) variants were further visualised with a *G*_*S*T_ scatter plot and the variants located in genetracts in PlasmoDB ^64^. Customised scrit with TopR and locuszoom package in R, were used applied to reveal details around differentiating loci. To extend the temporal analysis to include allele frequencies at intervening times over the period, previously obtained sequence data from 1984 and 2001 were analysed ^1^.

## Acknowledgements

We are grateful to current and previous members of staff at the MRC Unit in The Gambia and the London School of Hygiene and Tropical Medicine who at different times enabled the maintenance of research sample archives. We thank Lindsay Stewart for support with the DNA extraction processing, as well as Eleanor Drury, Victoria Simpson and Sonia Goncalves at the Wellcome Sanger Institute for support with processing of samples for sequencing.

This research was supported by a European and Developing Countries Clinical Trials Partnership (EDCTP) Senior Fellowship Plus award (TMA2019SFP-2843-EGSAT) and a Wellcome Sanger Institute Senior International Fellowship award (S4739-IF-AA-N) to AA-N and an MRC Project Grant (MR/S009760/1) to DJC.

