## Supplementary Fig. for "New insights on malaria parasite adaptation yielded by samples archived for more than 50 years"

### Died in April 2023

**A**

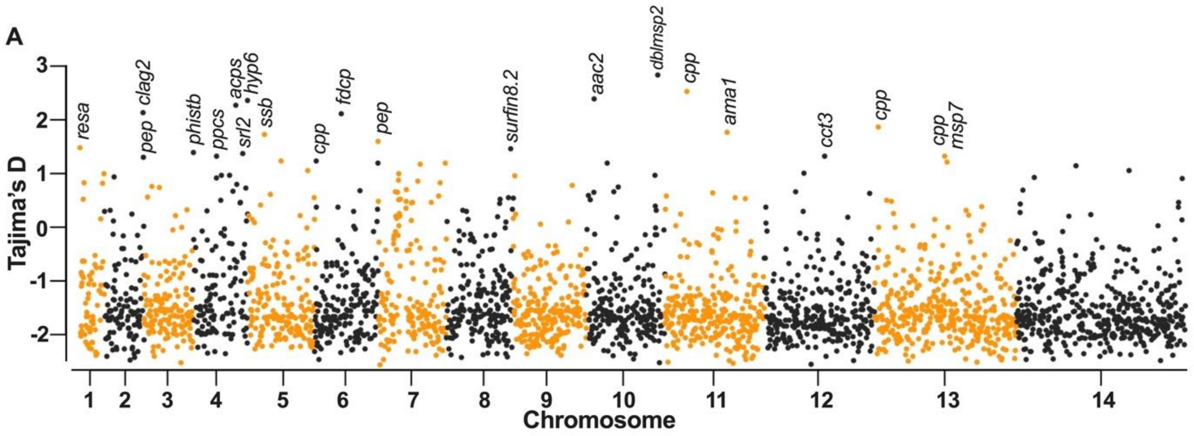


**B**

| Gene Symbol | Gene ID | Gene Description | N | S | Tajima's D |
|---|---|---|---|---|---|
| *resa* | PF3D7_0102200 | ring-infected erythrocyte surface antigen | 48 | 6 | 1.49 |
| *clag2* | PF3D7_0220800 | cytoadherence linked asexual protein 2 | 38 | 42 | 2.15 |
| *pep* | PF3D7_0221000 | Plasmodium exported protein, unknown function | 49 | 23 | 1.31 |
| *pfd80* | PF3D7_0401800 | Plasmodium exported protein, unknown function | 51 | 5 | 1.40 |
| *ppcs* | PF3D7_0412300 | phosphopantothenoylcysteine synthetase, putative | 49 | 10 | 1.33 |
| *acps* | PF3D7_0420200 | holo-[acyl-carrier-protein] synthase, putative | 50 | 15 | 2.29 |
| *sr12* | PF3D7_0422800 | serpentine receptor 12, putative | 46 | 10 | 1.38 |
| *hyp6* | PF3D7_0425100 | Plasmodium exported protein , unknown function | 47 | 27 | 2.38 |
| *ssb* | PF3D7_0508800 | single-stranded DNA-binding protein | 52 | 3 | 1.74 |
| *cpp* | PF3D7_0602700 | conserved Plasmodium protein, unknown function | 52 | 8 | 1.24 |
| *fdcp* | PF3D7_0614100 | filamin domain-containing protein, putative | 50 | 6 | 2.13 |
| *pep* | PF3D7_0701900 | Plasmodium exported protein, unknown function | 48 | 55 | 1.61 |
| *surfin8.2* | PF3D7_0830800 | surface-associated interspersed protein 8.2 | 26 | 141 | 1.47 |
| *aac2* | PF3D7_1004800 | ADP,ATP carrier protein 2 | 51 | 9 | 2.41 |
| *dblmsp2* | PF3D7_1036300 | duffy binding-like merozoite surface protein 2 | 45 | 23 | 2.86 |
| *cpp* | PF3D7_1113600 | conserved Plasmodium protein, unknown function | 50 | 5 | 2.55 |
| *ama1* | PF3D7_1133400 | apical membrane antigen 1 | 43 | 45 | 1.78 |
| *cct3* | PF3D7_1229500 | T-complex protein 1 subunit gamma | 54 | 4 | 1.33 |
| *cpp* | PF3D7_1302900 | conserved protein, unknown function | 47 | 17 | 1.88 |
| *msp7* | PF3D7_1335100 | merozoite surface protein 7 | 45 | 21 | 1.33 |

**Supplementary Fig 1**. Genome-wide scan for *P. falciparum* genes with more intermediate allele frequencies than expected indicating balancing selection in the sampled population from 1966 – 1971 in The Gambia. **A.** Tajima's D values are plotted for each genes containing at least 3 SNPs, and the genes with the top 20 highest values are labelled as these indicate the most exceptional patterns of intermediate allele frequencies. **B.** Description and summary of diversity for the genes having the top 20 values of Tajima's D index. N indicates the number of samples with sequences contributing to the analysis for each gene. S is the number of segregating sites.

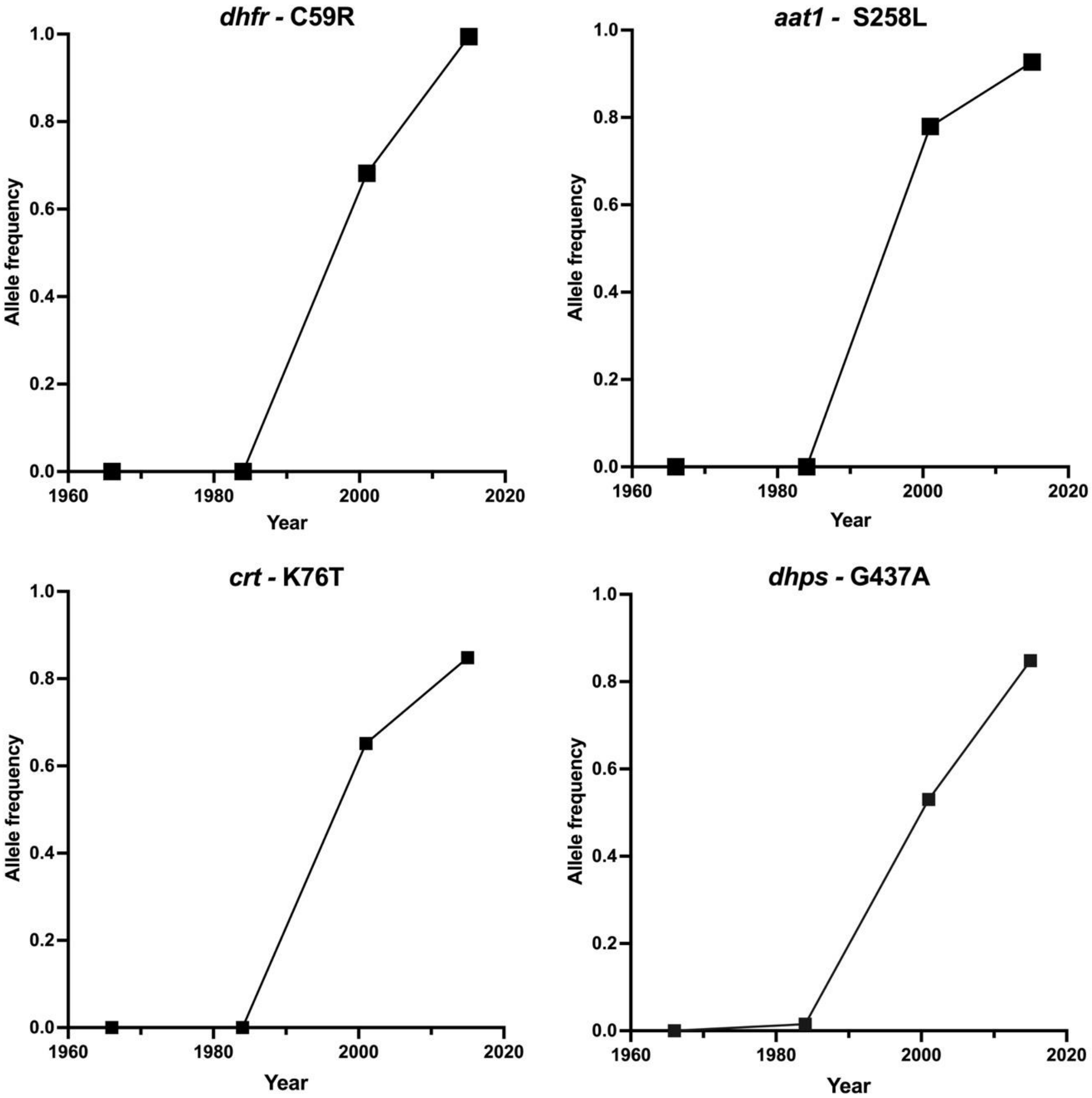


**Supplementary Fig. 2**. Temporal allele frequency changes over a period of almost 50 years in The Gambia for known drug resistance associated mutations in dihydrofolate synthetase (*dhfr*), amino acid transporter (*aat1*), chloroquine resistance transporter (*crt*) and dihydropteroate synthetase (*dhps*). A full list of non-synonymous SNPs with significant allele frequency change over time is presented in Supplementary Table S9.

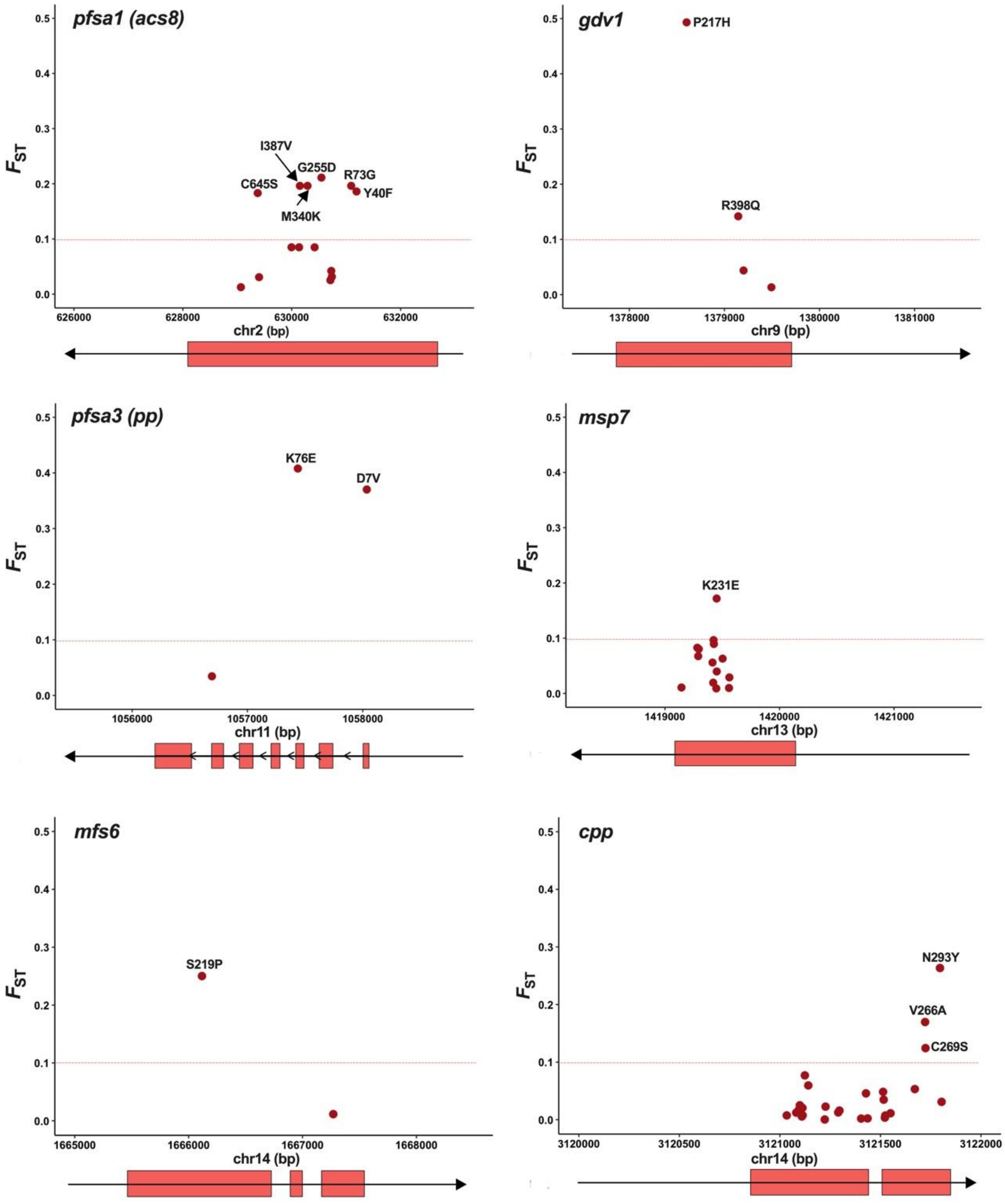


**Supplementary Fig. 3.** Graphical zoom in to individual *P. falciparum* gene loci with SNPs having highly differentiated frequencies ($F_{ST}$ fixation indices > 0.1) in The Gambia in a comparison between 1966-1971 and 2015. Each panel presents $F_{ST}$ values for SNPs plotted against their chromosomal position (chr) in the coding sequences (labelled with gene symbol in the top left). The protein position for amino acid alleles at non-synonymous variants with $F_{ST}$ > 0.1 are indicated. Below each $F_{ST}$ plot is a scheme showing the gene exons (red blocks), and the direction of transcription in relation to the genomic coordinates shown by arrow heads. The gene symbols represent: *pfsa1* = *P. falciparum* sickle associate locus 1 coding for acetyl coenzyme A synthetase 8 (*acs8*), *gdv1* = gametocyte development protein1, *pfsa3* = *P. falciparum* sickle associate locus 1 coding for a protein phosphatase, putative (*pp*), *msp7* = merozoite surface protein 7, *mfs6 =* major facilitator superfamily domain-containing protein, putative, and *cpp* = conserved *Plasmodium* protein.

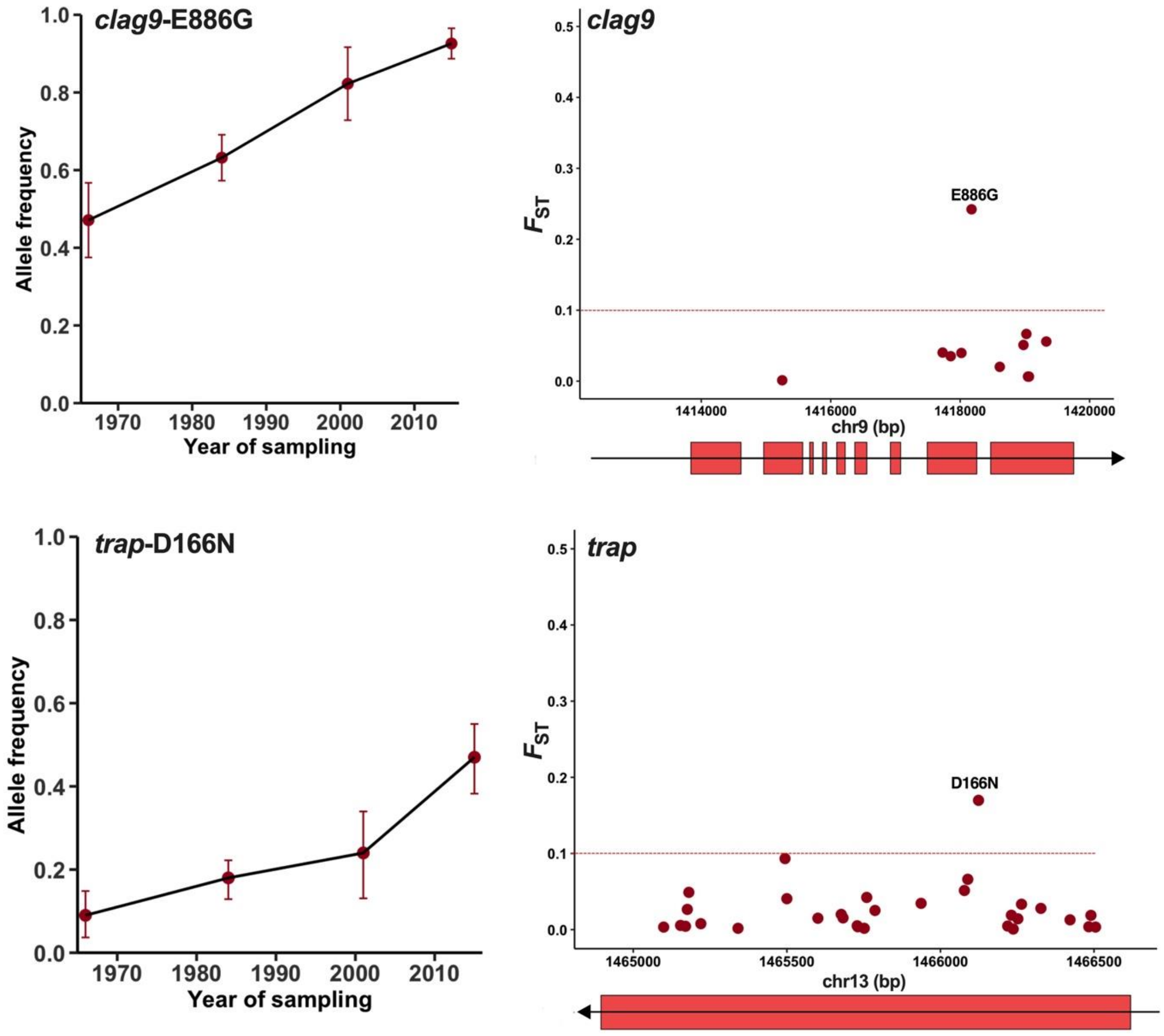


**Supplementary Fig. 4.** Allele frequency changes over time for SNPs in two genes closely linked with others showing significant changes in Fig. 4. The top row shows allele frequencies at different times from 1966-71 (plotted by with the median sample year 1968) onwards for variant E886G in *clag9* adjacent to a plot of the $F_{ST}$ values of all coding SNPs in the gene on chromosome 9. Exons depicted as red bars and arrows show the coding frame. The second row shows equivalent plots for the *trap* gene on chromosome 13, with allele frequencies showing frequency changes of the most temporally differentiated variant D166N.
